# The Ribosome Assembly Factor Reh1 is Released from the Polypeptide Exit Tunnel in the Pioneer Round of Translation

**DOI:** 10.1101/2023.10.23.563604

**Authors:** Sharmishtha Musalgaonkar, James Yelland, Ruta Chitale, Shilpa Rao, Hakan Ozadam, Can Cenik, David Taylor, Arlen Johnson

**Author notes:** Corresponding author. (AWJ). Basic Sciences Division and Computational Biology Section of the Public Health Sciences Division, Fred Hutchinson Cancer Center, Seattle, WA.

## Abstract

Assembly of functional ribosomal subunits and successfully delivering them to the translating pool is a prerequisite for protein synthesis and cell growth. In *S. cerevisiae,* the ribosome assembly factor Reh1 binds to pre-60S subunits at a late stage during their cytoplasmic maturation. Previous work shows that the C-terminus of Reh1 inserts into the polypeptide exit tunnel (PET) of the pre-60S subunit. Unlike canonical assembly factors, which associate exclusively with pre-60S subunits, we observed that Reh1 sediments with polysomes in addition to free 60S subunits. We therefore investigated the intriguing possibility that Reh1 remains associated with 60S subunits after the release of the anti-association factor Tif6 and after subunit joining. Here, we show that Reh1-bound nascent 60S subunits associate with 40S subunits to form actively translating ribosomes. Using selective ribosome profiling, we found that Reh1-bound ribosomes populate open reading frames near start codons. Reh1-bound ribosomes are also strongly enriched for initiator tRNA, indicating they are associated with early elongation events. Using single particle cryo-electron microscopy to image cycloheximide-arrested Reh1-bound 80S ribosomes, we found that Reh1-bound 80S contain A site peptidyl tRNA, P site tRNA and eIF5A indicating that Reh1 does not dissociate from 60S until early stages of translation elongation. We propose that Reh1 is displaced by the elongating peptide chain. These results identify Reh1 as the last assembly factor released from the nascent 60S subunit during its pioneer round of translation.

## The 60S cytoplasmic maturation pathway

In eukaryotic cells, ribosome synthesis begins co-transcriptionally on the nascent rDNA transcript in the nucleolus. Early events of rRNA processing and RNA folding in the nucleolus culminate in the release of pre-40S and pre-60S subunits into the nucleoplasm, from which they are independently exported to the cytoplasm (see ^1–4^ for recent reviews). Although the nascent subunits are largely assembled prior to nuclear export, they must undergo key cytoplasmic assembly events to become functional ribosomal subunits ^5,6^. These cytoplasmic assembly events include completion of the peptidyl transfer center. In addition to final assembly events, an entourage of assembly factors that block critical ligand binding sites on the ribosomal subunits must be removed to allow the ribosome to function in translation^7^. Notably, the nuclear export adapter Nmd3 occupies the P and E sites, Tif6 binds to the joining face, thereby preventing association with a small subunit, and the GTPase Nog1, which occupies the A site, also extends around the subunit and inserts into the polypeptide exit tunnel (PET)^8^.

In the cytoplasm, maturation of the pre-60S complex follows distinct hierarchical pathways^9^, which converge on a pre-60S intermediate bound by the ribosome assembly factors Nmd3, Tif6 and Reh1^10^. One branch of this pathway involves protein exchanges at the PET. ^1112^Because the C-terminus of Nog1 is intertwined with Rlp24, the release of Rlp24 is thought to simultaneously extract Nog1 from the PET^11^. Subsequently, Rei1 binds to the subunit and like Nog1, the C-terminus of Rei1 inserts into the exit tunnel^12^. The mechanism by which Rei1 is removed is not clearly understood, although it may be released in conjunction with Arx1 by the Hsp70 Ssa1/2^12–14^. In yeast, Rei1 has a paralog, Reh1, which replaces Rei1 at the time of Arx1 release^5^. Reh1 and Rei1 are functionally redundant^15^ and, as with Rei1, the C-terminus of Reh1 inserts into the exit tunnel^5,10^. In parallel with events at the exit tunnel, the peptidyl transferase center is completed by the insertion of Rpl10 (uL16) ^5,6^. Accommodation of Rpl10 triggers the release of Nmd3 from the P and E sites, allowing the P site ligand Sdo1 to bind. Sdo1 activates the GTPase Efl1 to release Tif6 from the nascent subunit^16^, thereby licensing the subunit for translation. We previously showed that Efl1 is sensitive to perturbations in the P site^17^, and we proposed that the Efl1-dependent release of Tif6 represents a “test drive” of the subunit, which simultaneously assesses the assembly of the P stalk and the integrity of the P site.

In our current understanding of 60S maturation, the release of Tif6 is thought to be the last step, releasing a fully mature ribosome, free of assembly factors, into the translating pool. However, the timing and mechanism of release of Reh1 have not been examined. While the C-terminus of Reh1 occupies the exit tunnel, the N-terminus of Reh1 is thought to interact directly with Tif6^5^, leading to the notion that Reh1 is released in conjunction with Tif6. This idea is supported by structural analysis, which identified a late maturation intermediate containing Tif6 and Reh1, but lacking Nmd3 and Lsg1^5^. Here, we show that Reh1 is released not only after Tif6, but after translation initiation. Our results suggest that Reh1 is released at an early stage of translation elongation, possibly triggered by the nascent polypeptide. Our findings identify Reh1 as a non-canonical ribosome assembly factor that remains associated with the nascent subunit through translation initiation and uniquely marks the pioneer round of translation for the newly made 60S subunit.

## Results

### Reh1 sediments with free 60S subunits and polysomes

Mass spectrometric and cryo-EM analyses have identified Reh1 as a late 60S assembly factor that joins the pre-60S after the release of its paralog, Rei1, in the cytoplasm ^5,10,18^ and all Reh1-containing particles also contain Tif6. As the release of Tif6 licenses pre-60S for translation by relieving the block in joining to 40S (Figure 1A), the release of Tif6 has been assumed to be the last step of 60S maturation. However, the timing of Reh1 release relative to Tif6 release remains unknown. We monitored the sedimentation of Reh1 in sucrose density gradients to ask if it behaved as other canonical pre-60S factors which sediment exclusively with pre-60S. In contrast to Nmd3, which was present exclusively on free 60S subunits, Reh1 was present not only on free 60S subunits but also in polysome-containing fractions (Figure 1B). This result is consistent with previously published work^15^ and suggests that Reh1 persists on 60S subunits after the release of Tif6 and after subunit joining.

**Figure 1.**
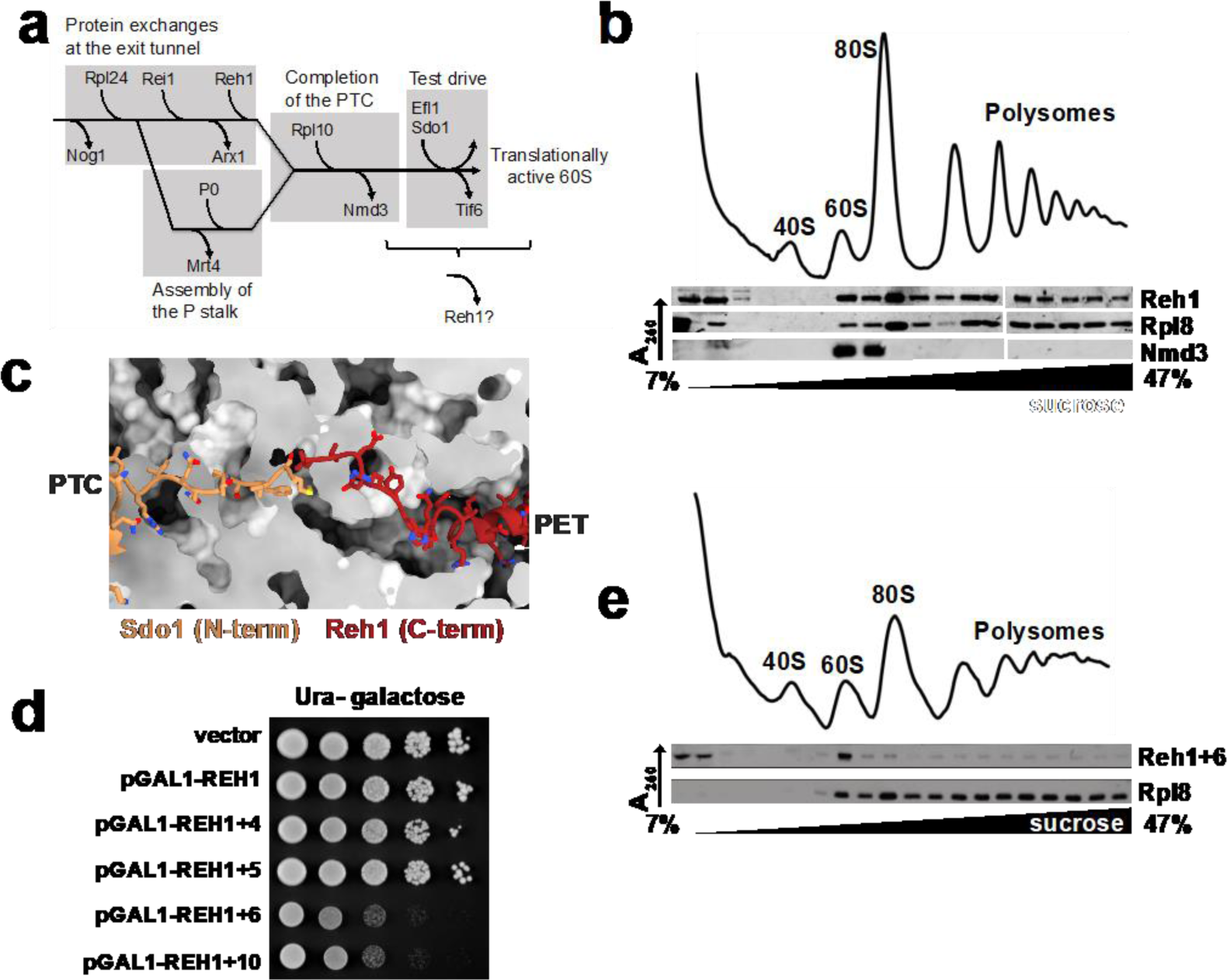
Reh1 remains associated with 60S during the transition from biogenesis to translation. *Panel A:* Simplified pathway of cytoplasmic 60S maturation. The major events at the exit tunnel, assembly of the P-stalk, completion of the peptidyl transferase center (PTC) and the test drive to license the subunit are shown. *Panel B:* Reh1 sediments into polysomes. Extract of WT cells (AJY4408) expressing 3xmyc-tagged Reh1 (pAJ4763) was separated by sucrose density gradient sedimentation. UV trace monitoring A_260_ is shown. Fractions were analyzed by western blotting for the presence of Reh1, Rpl8 and Nmd3. *Panel C*: Cartoon depicting the expected relationship of the N-terminus of Sdo1 (human SBDS, PDB 5AN9^54^) and the C-terminus of yeast Reh1 (PDB 6QTZ^5^) in the PET of the 60S subunit. *Panel D*: REH1+6 is dominant negative. Ten-fold serial dilutions of wild-type yeast (BY4741) containing vector (pRS416), or galactose-inducible WT Reh1 (pAJ4763) or REH with the indicated number of amino acids from the N-terminus of Sdo1 fused to the C-terminus of REH1) plated on galactose-containing media. *Panel E:* Reh1+6 does not enter polysomes. Extract of cells expressing Reh1+6 (pAJ4764) was separated by sucrose density gradient sedimentation. UV trace and the sedimentation of Reh1+6 and Rpl8 are shown.

### Reh1 progresses from pre-60S into polysomes

Our results suggest that Reh1 remains associated with 60S subunits as they enter the translation cycle. However, it was also possible that Reh1 enters polysomes by binding to recycling mature subunits. To ask if Reh1 enters polysomes exclusively through the biogenesis pathway, we designed an Reh1 mutant that would block the progression of nascent 60S into the pool of actively translating ribosomes. We reasoned that extending the C-terminus of Reh1, which inserts into the polypeptide exit tunnel, would block Sdo1 binding and arrest 60S maturation at the point of Tif6 release. To this end, we extended the C-terminus of Reh1 with residues from the N-terminus of Sdo1, which inserts into the exit tunnel from the joining face (Figure 1C). We made a series of Reh1 variants in which we extended the C-terminus with four, five, six and ten residues from the N-terminus of Sdo1.

Overexpression of variants with four or five residue extensions had no adverse effect on cells but variants extended by six (Reh1+6) or ten residues had a potent dominant negative effect when overexpressed (Figure 1D). As predicted, the sedimentation of Reh1+6 was restricted to pre-60S (Figure 1E). The absence of Reh1+6 from polysome fractions strongly suggests that this protein does not bind to recycling 60S, supporting our conclusion that WT Reh1 enters polysomes through the biogenesis pathway.

### A dominant mutant of Reh1 blocks the release of Nmd3 and Tif6

We designed Reh1+6 expecting that it would block Sdo1-stimulated release of Tif6 by the GTPase Efl1^16^. To directly test if Reh1+6 inhibited Efl1 function, we assayed the effect of Reh1+6 on the Sdo1-dependent activation of Efl1 GTPase activity. Efl1 GTPase activity is activated by Sdo1 in the presence of 60S subunits (Figure 2A). Addition of increasing amounts of Reh1 did not significantly affect Efl1 activity. However, Reh1+6 had a dose dependent inhibitory effect on Efl1 (Figure 2A).

**Figure 2.**
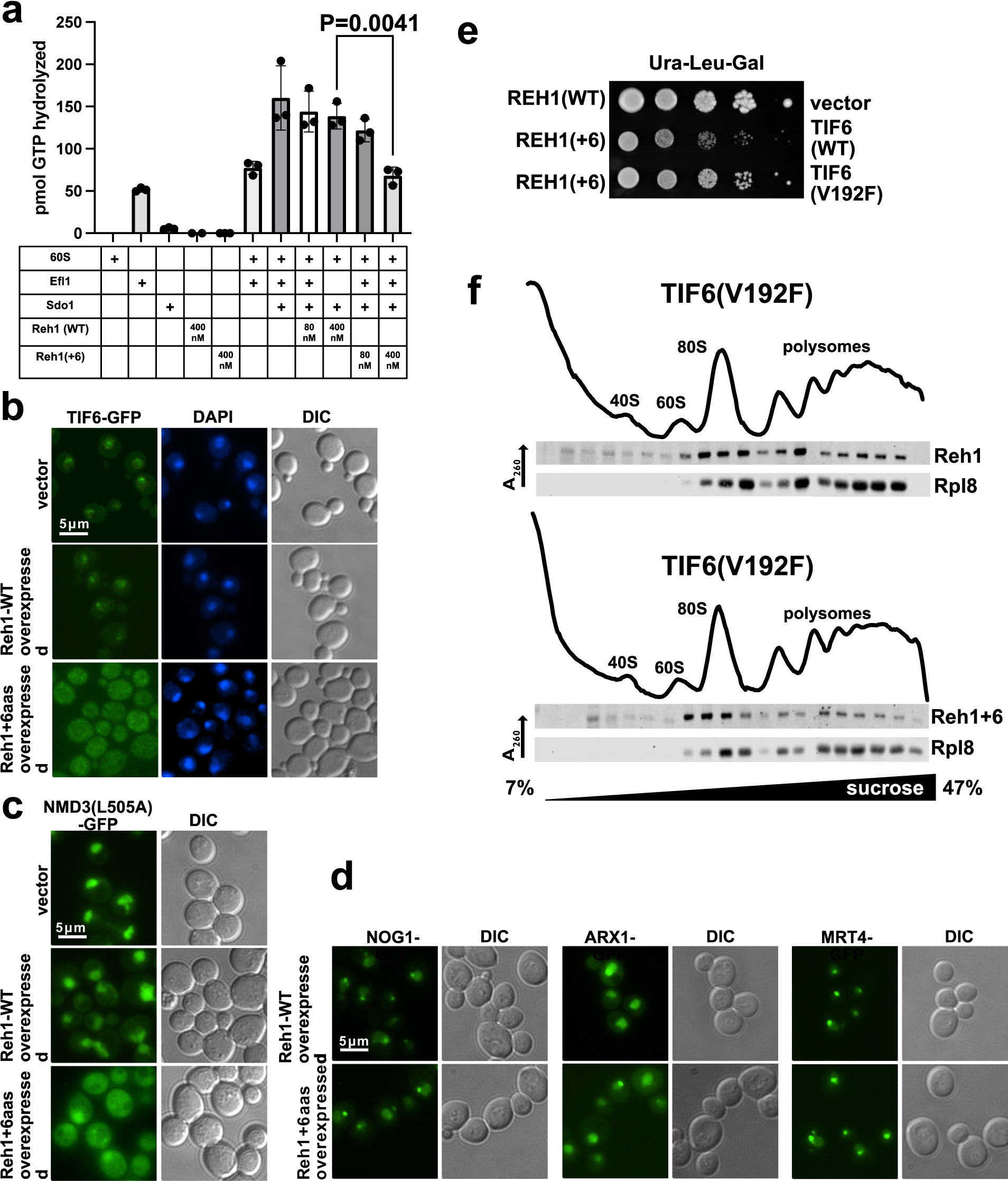
Reh1+6 blocks Sdo1 function. *Panel A:* Reh1+6 inhibits Sdo1-dependent GTPase activity of Efl1. Sdo1 stimulated, 60S-dependent Efl1 GTPase activity was monitored by the production of GDP in reactions containing the indicated combinations of 80nM 60S subunits, 100nM Sdo1, 60nM Efl1 and 80nM or 400nM Reh1 (WT or mutant). All experiments were done in Triplicate and significance was evaluated with a two-tailed t test. *Panel B:* Fluorescence microscopy of TIF6-GFP expressing cells (AJY4049) containing vector or expressing Reh1 WT (pAJ4751) or Reh1+6 (pAJ4157). DAPI staining for nuclei and DIC images are shown for reference. *Panel C:* Similar to panel A except the Nmd3-GFP expressing strain AJY4050 was used. *Panel D*: Reh1+6 does not block recycling of the upstream factors Nog1, Mrt4 and Arx1. Similar to panel A except that strains were AJY4659 (NOG1-GFP), AJY4661 (ARX1-GFP) and AJY4660 (MRT4-GFP) and only DIC is shown for reference. *Panel E:* The dominant negative effect of Reh1+6 is suppressed by TIF6(V192F). Ten-fold serial dilutions of wild-type yeast (BY4741) containing vectors expressing the indicated factors were plated on galactose-containing medium. Vectors used were: REH1 (WT) pAJ4751, REH1+6 (pAJ4157), vector (pRS415), TIF6 (WT) pAJ2846, TIF6-V192F pAJ2249. *Panel F:* Reh1+6 enters polysomes when suppressed by TIF6(V192F). Extracts of TIF6-V192F mutant cells (AJY4659) expressing REH1 WT (pAJ4751) or REH1+6 (pAJ4157) were separated by sucrose density gradient sedimentation. UV trace monitoring rRNA is shown. Fractions were analyzed by western blotting for the presence of Reh1 and Rpl8.

To determine at which step Reh1+6 arrested the cytoplasmic maturation pathway *in vivo*, we examined the cellular localization of assembly factors that shuttle between the nucleus and cytoplasm ^9^. Overexpression of the dominant negative Reh1+6, but not WT Reh1, strongly inhibited nuclear recycling of Tif6 (Figure 2B). To monitor the recycling of Nmd3, which is cytoplasmic at steady state, we employed the Nmd3-L505A mutant, which we previously found to have a reduced rate of nuclear export, shifting the steady state distribution of Nmd3 to the nucleus^19^. Overexpression of Reh1+6 also strongly inhibited the nuclear recycling of Nmd3-L505A (Figure 2C). In contrast, overexpression of Reh1+6 did not block the recycling of Nog1, Arx1 or Mrt4, which are released from pre-60S in the cytoplasm upstream of the release of Nmd3 and Tif6 (Figure 2D) ^9^. Thus, Reh1+6 specifically blocks a late step in 60S assembly.

Sdo1 and Efl1 are dispensable in the presence of *TIF6* mutants that weaken the affinity of Tif6 for pre-60S^20^. We tested if one such mutant, TIF6-V192F, could bypass the dominant negative effect of Reh1+6. Indeed, co-expression of Reh1+6 with TIF6-V192F but not WT TIF6 largely relieved the dominant negative growth effect of Reh1+6 (Figure 2E). Remarkably, TIF6-V192F also allowed Reh1+6-containing 60S subunits to enter polysomes (Figure 2F). These results show that Reh1+6 can enter polysomes, but only when the biogenesis arrest is alleviated by mutation of *TIF6*. Furthermore, these results indicate that Reh1 is released after Tif6. We conclude that Reh1 is the last biogenesis factor to be released from the 60S subunit, and that its release occurs after subunit joining.

### The C-terminus of Reh1 is critical for function and for binding to 60S subunits

Structural studies of Reh1 on pre-60S have shown that the extreme C-terminus occupies the PET (Figure 2 S1A), while the N-terminus of the protein interacts with Rpl24 and with Tif6 on the joining face^21,22^. Thus, after Tif6 is released, Reh1 may be tethered to the ribosome primarily through the interaction of its C-terminus with the PET. To ask if the C-terminus contributes to Reh1 binding, we expressed C-terminal truncations *in vivo* and assayed their sedimentation with ribosomes in sucrose density gradients. We generated two truncations that removed 6 or 50 residues from the C-terminus, based on the structure of Reh1 in the PET (Figure 2 S1A). The 6 amino acid deletion removed residues extending beyond the uL22-uL4 constriction point in the PET, whereas the 50 amino acid deletion removed the entirety of the C-terminus that occupies the exit tunnel. We tested the ability of these truncations to complement loss of REH1. Because REH1 is partially redundant with REI1^15^, we tested complementation in a reh1Δ rei1Δ double mutant. However, because Rei1 is needed for the removal of Arx1 from pre-60S^21,22^, we also deleted ARX1 from this strain to suppress the loss of REI1^21^. We found that deletion of either 6 or 50 amino acids from the C-terminus of Reh1 rendered the protein nonfunctional (Figure 2 S1B). However, the two truncations differed in their ability to bind to ribosomes: Reh1Δ6 retained ribosome binding, whereas Reh1Δ50 did not (Figure 2 S1C). These results demonstrate that the C-terminal 50 amino acids of Reh1 are important for its association with the ribosome. It is surprising that Reh1Δ6 did not complement the loss of functional Reh1, as Reh1Δ6 still binds the ribosome. This result suggests that the C-terminus of Reh1 contributes an additional but unidentified function.

### Selective ribosome profiling of Reh1-bound ribosomes

Thus far, our results indicate that Reh1 remains associated with nascent 60S subunits after subunits join during translation initiation. To gain a more complete understanding of when Reh1 is released during the translation cycle, we used selective ribosome profiling to determine where Reh1-containing ribosomes are positioned on mRNAs (Figure 3A).

**Figure 3.**
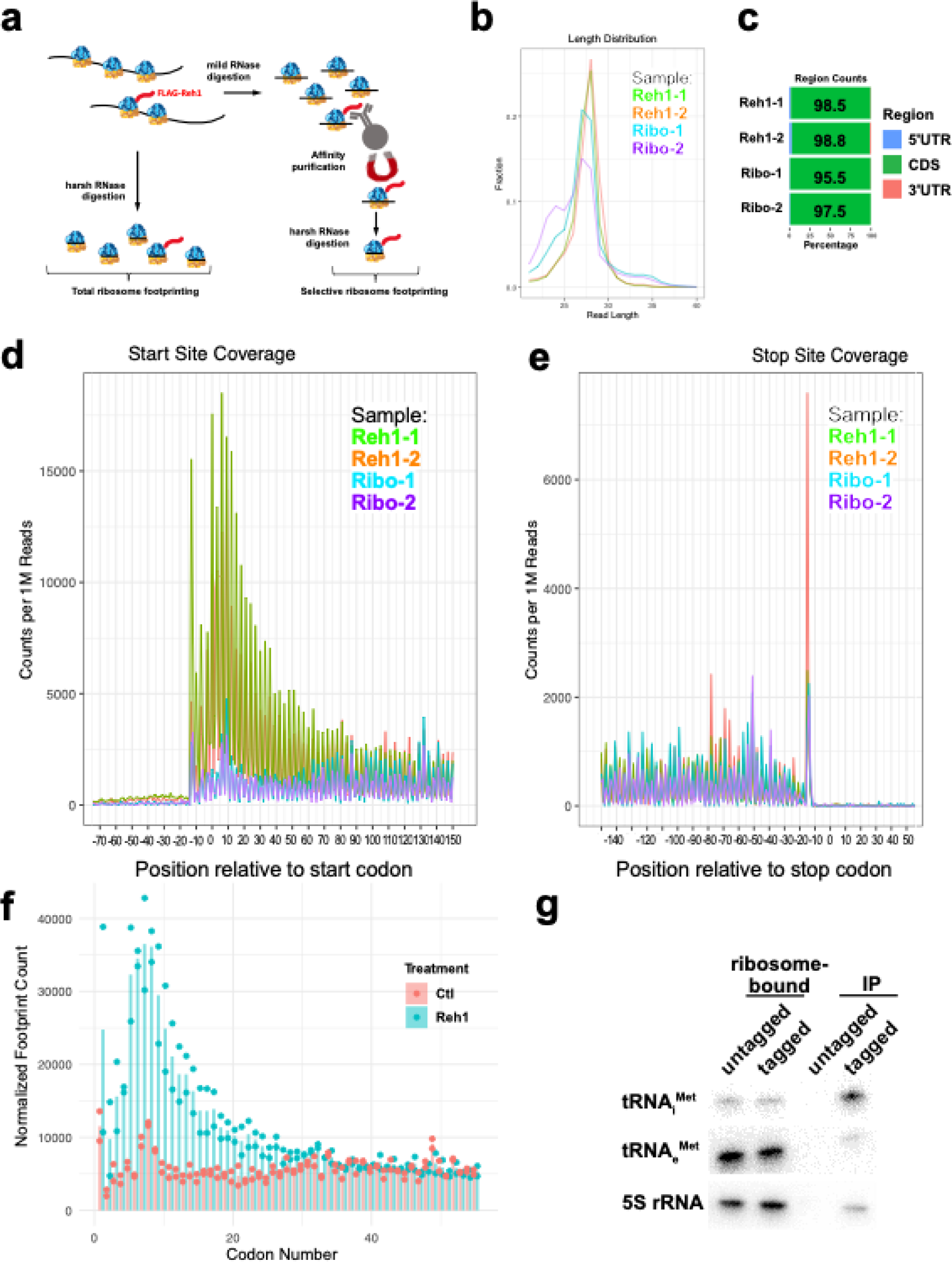
Reh1 is associated with translating ribosomes near the start codon. *Panel A*: Schematic of selective ribosome profiling. *Panel B*: The fraction of specific read lengths among the total mapped reads is plotted. *Panel C:* The percentage of ribosome profiling reads across 5’-UTR, CDS, 3’-UTR regions. *Panel D:* Metagene analysis of footprints mapping near the start codons. *Panel E*: Metagene analysis of footprints mapping near the stop codons. *Panel F*: Distribution of reads surrounding start codons were aggregated by codon number and the mean number of reads (bar) along with individual replicates (points) were plotted. *Panel G*: Enrichment of initiator tRNA ^Met^ in Reh1-bound ribosomes. Reh1 was immunoprecipitated from RNaseI-treated extracts of cells expressing untagged or Protein A-tagged Reh1. In parallel, bulk ribosomes were sedimented through sucrose cushions to isolate ribosome-associated tRNAs. RNAs were separated on a polyacrylamide gel, transferred to membrane and probed for 5S rRNA, methionine elongator tRNA (tRNA_e_^Met^) and initiator tRNA (tRNA ^Met^).

The footprints from Reh1-bound ribosomes that mapped to mRNAs displayed a length distribution centered on 28 nt (Figure 3B), and the vast majority (>95%) originated from the coding regions (Figure 3C). Similar results were obtained with bulk ribosome profiling suggesting the high quality of these datasets (Figure 3C). Both Reh1-bound and bulk ribosome footprints displayed clear 3 nucleotide periodicity (Figure 3D), indicative of translating ribosomes.

Excitingly, analysis of the Reh1-bound ribosome footprints showed a strong enrichment within approximately the first 10 codons of open reading frames (ORFs), compared to bulk ribosome footprints, which were uniformly distributed across coding regions (Figure 3D). In contrast, metagene analysis did not reveal consistent enrichment of selected Reh1 footprints in the region preceding the stop codons (Figure 3E). The strong enrichment of Reh1-bound ribosome footprints at the 5’-end of ORFs is consistent with a model in which Reh1 is released in early rounds of translation as the growing nascent polypeptide encounters the C-terminus of Reh1. Based on structural models of the C-terminus of Reh1, between 10-15 residues of extended polypeptide can be accommodated between the acceptor stem of tRNA and the constriction site of the PET, where it is likely that nascent polypeptide and the C-terminus of Reh1 would be incompatible. This distance correlates well with the enrichment of Reh1-bound ribosome footprints within the first 10 codons (Figure 3F), corresponding to ribosomes containing nascent polypeptides of 2-10 amino acids.

### Reh1 particles are enriched for initiator tRNA

The notion that Reh1 may be evicted by the nascent polypeptide during early stages of translation together with the observation that Reh1 is enriched on mRNAs near the start codon, prompted us to ask if Reh1-bound ribosomes are enriched for initiator tRNA (tRNA_i_ ^Met^). We compared the relative enrichment of initiator tRNA_i_^Met^ and elongator tRNA_e_^Met^ in Reh1-bound ribosomes to total ribosome pool. We observed a strong (∼10-fold) enrichment of tRNA_e_^Met^ relative to tRNA_e_^Met^ in the Reh1-bound sample (Figure 3G). The enrichment of tRNA_i_ ^Met^ was also greater than the enrichment of 5S rRNA, as expected if Reh1-bound ribosomes are enriched for tRNA_i_ ^Met^, whereas all ribosomes, free and Reh1-bound contain 5S. Both tRNAs and 5S rRNA were virtually undetectable in the untagged negative control, showing that the RNA species were specifically associated with Reh1. Reh1-containing ribosomes were also enriched for tRNA_i_^Met^ relative to tRNA_e_^Met^ (∼5-fold) and tRNA ^Arg^ when compared to the total input RNA before RNase I digestion (Figure 3 S1A). Enrichment for tRNA_i_^Met^ relative to tRNA_e_^Met^ (∼10-fold) was also seen when compared to RNAse-treated extracts (Figure 3 S1B). Although we do not know the true stoichiometry of initiator tRNA in the Reh1 complexes without a standard that is fully occupied with tRNA_i_^Met^, our results show that Reh1-associated ribosomes are strongly enriched for initiating 80S ribosomes, consistent with our ribosome profiling analyses.

### Cryo-EM structure of the Reh1-80S complex

To gain structural insight into the nature of the Reh1-bound 80S complex, we determined the 3.1 Å cryo-EM structure of immunoprecipitated Reh1-3xFLAG bound to the translating ribosome. We used a focused 3D classification strategy to separate 80S complexes by occupancy of the A site and E site, resolving four distinct ribosomal complexes (Supplemental figure 4S1B). Two complexes were occupied by A and P site tRNAs and mRNA with eIF5A (Reh1-80S-tRNA-eIF5A, Figure 4A) and without eIF5A (Reh1-80S-tRNA, Supplemental figure 4S2A). One complex lacked tRNAs but contained eIF5A and the dormancy factor Stm1 (Supplemental Figures 4S2B), which has previously been observed in both vacant and eIF5A-containing ribosome structures^23,24^. The fourth complex lacked tRNAs and translation factors. Our structures reveal a translating 80S complex in which the C-terminal 57 amino acids of Reh1 occupy nearly the entire length of the exit tunnel (Figure 4A). In these structures, the C-terminus of Reh1 adopts a conformation nearly identical to that observed in the exit tunnel of pre-60S subunits^5,10^. The remaining 375 amino acids of Reh1 were not observed in any of our structures, suggesting that the C-terminus of Reh1 forms the only stable connection between Reh1 and the translating 80S.

**Figure 4.**
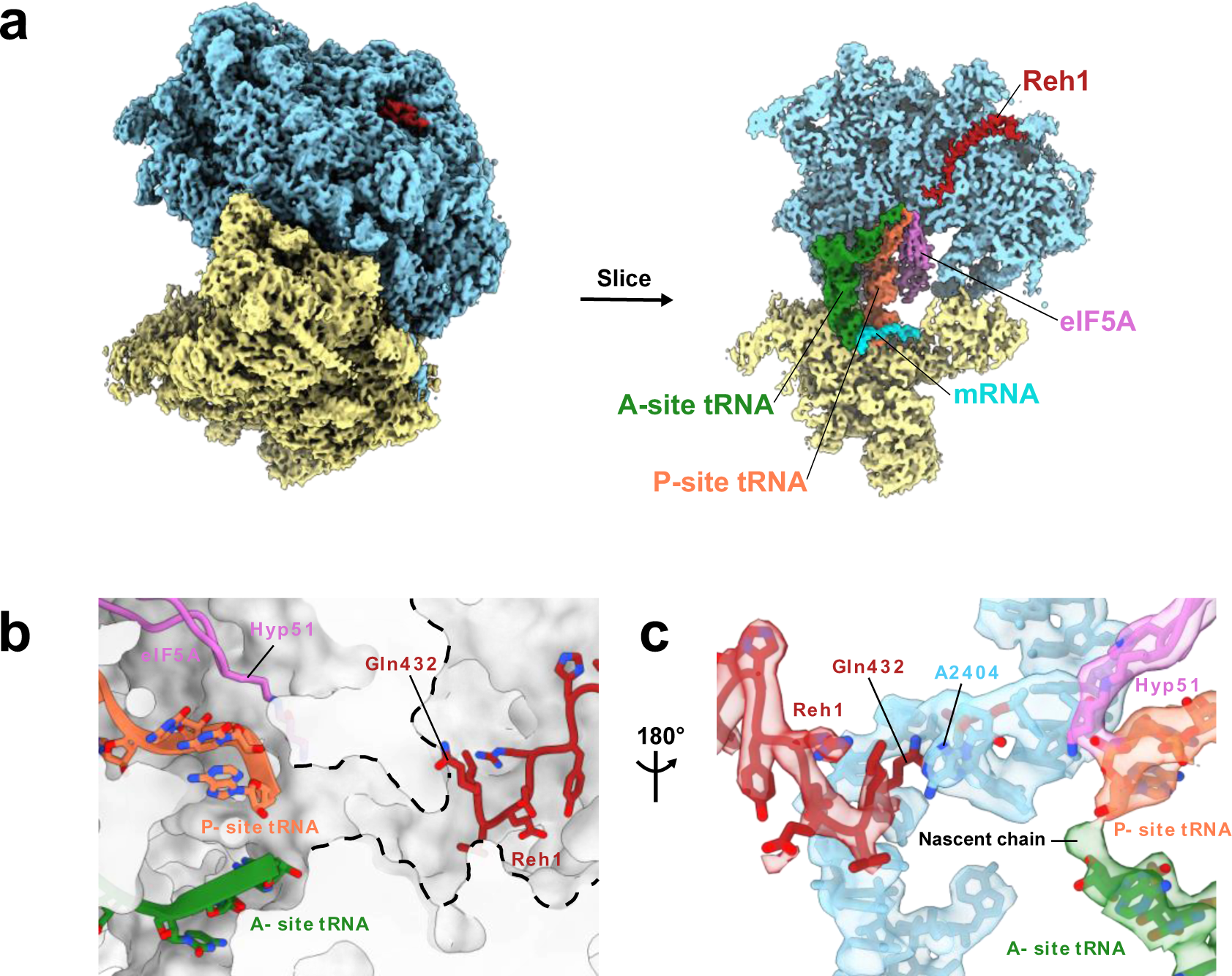
Cryo-EM structure of Reh1-bound 80S. *Panel A:* Cryo-EM map (left) and cross-section (right) of the eIF5A-containing Reh1-80S structure with Reh1 (red), A site tRNA (green), P site tRNA (orange), mRNA (turquoise), and eIF5A (purple). Ligands are surrounded by the large ribosomal subunit (blue) and small ribosomal subunit (yellow). *Panel B*: Cartoon representation of eIF5A, P site tRNA, and A site tRNA in the active site of Reh1-80S-tRNA-eIF5A. The PET is delineated with a dashed line. *Panel C:* Detailed view of cryo-EM map and model of Reh1-80S-tRNA-eIF5A, showing Reh1 interaction with constrictor site base A2404. Shown are Reh1 (red), A site tRNA (green), P site tRNA (orange), eIF5A (purple), and residues 2393-2407 of 25S rRNA (blue).

Nearly half of Reh1-containing 80S particles contained mRNA and tRNAs in the A and P sites. Within 42% of these 80S complexes, the essential translation factor eIF5A was also present in the E site. The unexpected observation of eIF5A reinforces the notion that Reh1-80S structures represent actively translating ribosomes captured at a relatively slow step of elongation. Our finding also raises the possibility that eIF5A is needed for, or accelerates, dissociation of Reh1 during the first rounds of translation. However, the structure of the large subunit precludes eIF5A from making direct contact with Reh1. The critical modified amino acid of eIF5A, hypusine 51 (Hyp51), is inserted deep into the PTC E site, in proximity to the 3’ end of P site tRNA (Figure 4B). The C-terminal residue of Reh1, glutamine 432 (Gln432), forms a stacking interaction with adenosine 2404 (A2404) of the 25S rRNA (Figure 4C): a component of the PET wall, located approximately 14 Å from the 3’ hydroxyl of A site tRNA. Importantly, A2404 separates the PET from the PTC. Therefore, it is likely that nascent peptide chain extension directly evicts Reh1 from the 80S, perhaps by disrupting stable interactions between the C-terminus of Reh1 and the 25S rRNA. In our structures of 80S containing tRNA, the ribosomes are stalled in the pre-rotated state by cycloheximide (Supplemental figure 4S2C) which was used to prevent ribosome runoff and displacement of Reh1. In these maps, a volume corresponding to ∼2 amino acids is seen to be continuous with the A site tRNA (Figure 4C). This observation suggests that further extension of the nascent chain is not compatible with a stably bound Reh1, a finding supported by our ribosome profiling results showing rapid depletion of Reh1-bound footprints after just a few rounds of elongation.

The presence of eIF5A is accompanied by closure of the L1 stalk, a shift of P site tRNA towards the A site, and slight closure of the PET around a fulcrum formed by the constriction site (Supplementary Video 1). These large-scale motions are potentially important for general eIF5A-catalyzed peptide bond formation. They could also be important for the release of Reh1, by decreasing the free energy of disrupting stabilizing interactions with the PET.

Our structures of tRNA-containing cycloheximide-arrested ribosomes also revealed that the N-terminus of eS25 (Rps25) intercalates between the anticodon stem loop (ASL) helices of A and P site tRNAs where a series of evolutionarily-conserved lysines make backbone contacts with both tRNAs (Supplemental figure 4S2D-F). The N-terminus of eS25 is conserved throughout eukaryotes, and has been reported to interact with the IRES of Cricket paralysis virus during translation initiation^25^. An interaction of eS25 with A and P site tRNAs has not been previously reported, but a similar interaction has been described for the C-terminus of uS19 (Rps15 in *S. cerevisiae*), which inserts between the A and P site tRNAs in cycloheximide-arrested elongating ribosomes from human cells ^26^ as well the filamentous fungus *N. crassa*^27^. It is tempting to speculate that the N-terminus of eS25 plays a specific role during the earliest stages of elongation, a state which we have captured with Reh1-bound ribosomes. Additional experimental work is required to test this intriguing possibility.

## Discussion

Maturation of the nascent 60S subunit in yeast has been thought to culminate with the release of the subunit anti-association factor Tif6 to license the subunit for translation^3^. Here, we have shown that the ultimate step of maturation is the release of the assembly factor Reh1 from the exit tunnel and that this step occurs after subunit joining at an early stage of translation elongation.

Previously work hinted at the possibility that Reh1 associates with translating ribosomes^15^. Our work conclusively shows this to be the case. Using selective ribosome profiling, we show that Reh1-containing ribosomes are strongly enriched at start codons and coding sequences immediately downstream of translation start sites. This conclusion is supported by the strong enrichment for initiator tRNA_i_^Met^ in Reh1-bound ribosomes. Finally, structures obtained by single-particle cryo-EM show that Reh1-bound ribosomes contain A and P site tRNAs engaged with mRNA and short nascent chains. In these structures, the C-terminus of Reh1 occupies the PET. As the presence of Reh1 in the PET is incompatible with the extension of the nascent polypeptide chain, we suggest that the growing nascent chain directly displaces Reh1 from the PET. Approximately half of the translating Reh1-bound ribosomes contained eIF5A in addition to A and P site tRNAs. It is notable that eIF5A was initially identified biochemically as a factor that promoted the first polypeptide bond in in vitro translation assays ^28–30^. Though it has been shown that eIF5A samples the E site of ribosomes throughout the length of elongation, and plays a particularly important role in the translation of certain codon pairs, eIF5A-containing ribosomes are enriched in the 5’-end of coding sequences^31^. These results, and those we present here, lead us to consider that eIF5A may play a role in facilitating the release of Reh1, either by accelerating the initial rounds of peptide bond formation or through a more complex allosteric mechanism.

It is remarkable that in eukaryotic cells, the PET is continuously occupied from the time of its initial formation during nucleolar steps in pre-60S assembly until the first translation event of the newly assembled subunit. During early stages of 60S assembly, the C-terminus of Nog1 scaffolds elements of the PET wall to stabilize the core pre-60S complex^32^ However, when Nog1 is released from the pre-60S in the cytoplasm, the structure of the PET is complete, and it is unlikely that the subsequent insertion of Rei1 followed by Reh1 plays a similar role in stabilizing the PET. Consequently, it is an open question what function these proteins contribute by continuing to occupy the exit tunnel after its assembly. One possibility is that these proteins prevent extraneous small molecules or peptides from lodging in the ribosome before it has initiated translation. Although most antibiotics target bacterial ribosomes, some can also act on eukaryotic ribosomes^33^. Small antimicrobial peptides which bind in the PET can also act on eukaryotic ribosomes^34^. Similarly, it is conceivable that endogenous, small (micro)peptides from the apparently ubiquitous translation of the genome could occupy the exit tunnel and stall translation^35^. In this scenario, Rei1 and Reh1 could keep the exit tunnel free of small molecules to ensure the unobstructed exit of the nascent polypeptide as the ribosome engages in translation.

Alternatively, Reh1 could surveil newly assembled ribosomes for function. Our results identify Reh1 as a factor that uniquely marks the 60S subunit in its pioneer round of translation; no other protein is known to associate exclusively with the nascent subunit to distinguish it from recycling mature subunits. If Reh1 is evicted by extension of a nascent polypeptide during a ribosome’s pioneer round of translation, the persistence of Reh1 on the ribosome could flag it as defective.

Although yeast contain the paralogous proteins Rei1 and Rh1, most eukaryotic species express a single Rei1-like protein. This is the case in human cells, which express only the Rei1-like protein ZNF622. The lack of an Reh1 homolog humans raises the question of whether ZNF622 fulfills the role of both Rei1 and Reh1, or Reh1 has acquired an additional function in yeast. The release of Reh1 in yeast is thought to be coincident with the release of Arx1 and precedes the release of Tif6. In human cells, the levels of ZNF622 depend on the ubiquitin E3 ligase HECTD1^36^. In the absence of HECTD1, ZNF622 accumulates on 60S subunits and causes the accumulation of eIF6 (Tif6) This result suggests that ZNF622 stabilizes eIF6 on pre-60S subunits, raising the possibility that ZNF622 is released after eIF6. If that is true, the release of ZNF622 would occur considerably later than that of its yeast counterpart Rei1, making it more similar to Reh1. While ZNF622 has not been reported to associate with translating ribosomes in human cells, this idea merits more direct analysis.

## Acknowledgements

This work was supported by NIH grants GM127127 (to A.W.J.) R35GM138348 (to D.W.T). and GM150667 (to CC), and Welch Foundation Research Grants F-1938 (to D.W.T) and F-2027-20230405 (to C.C.). C.C. is a CPRIT Scholar in Cancer Research supported by the Cancer Prevention and Research Institute of Texas grant RR180042. We thank D. Gottschling for pAOL47-04, J. Tesmer for pMALC2H10T, J. Dueber for yeast toolkit vectors and K.-Y. Lo for rabbit anti-Rpl8 antibody.

## Data and materials availability

Sequencing files for selective ribosome profiling experiments will be available at GEO (accession number GSEXXX). The structures of Reh1-80S-tRNA-eIF5A and Reh1-80S-tRNA have been deposited in the Electron Microscopy Data Bank as EMDB-xxxx and EMDB-yyyy, respectively. The corresponding molecular models of Reh1-80S-tRNA-eIF5A and Reh1-80S-tRNA have been deposited in the Protein Data Bank as PDB-zzzz and PDB-wwww, respectively.

---

**Table I.** Plasmids.

| Plasmid | Features | Source |
| --- | --- | --- |
| pRS415 | LEU2 CEN Ampr | <sup>37</sup> |
| pRS416 | URA3 CEN Ampr | <sup>37</sup> |
| pRS423 | HIS3 CEN Ampr | <sup>37</sup> |
| pAOL47-04 | pACT1-GAL4-EstrogenBD-VP16--NATMX<br>AmpR | Gottschling |
| pAJ2249 | TIF6-V192F LEU2 CEN Ampr | This work |
| pAJ2846 | TIF6 LEU2 CEN Ampr | <sup>38</sup> |
| pAJ4155 | PGAL1-REH1+4 URA3 CEN Ampr | This work |
| pAJ4156 | PGAL1-REH1+5 URA3 CEN Ampr | This work |

| Table I Plasmids |  |  |
| --- | --- | --- |
| Plasmid | Features | Source |
| pAJ4157 | PGAL1-REH1+6 URA3 CEN Ampr | This work |
| pAJ4612 | T7-MBP-HIS10-TEV-REH1 (WT) Ampr | This work |
| pAJ4613 | T7-MBP-HIS10-TEV-REH1 (+6) Ampr | This work |
| pAJ4618 | pGAL1-REH1 LEU2 CEN Ampr | This work |
| pAJ4619 | pGAL1-REH1+10 LEU2 CEN Ampr | This work |
| pAJ4621 | pGAL1-ProtA-TEV-FLAG-REH1 LEU2 CEN Ampr | This work |
| pAJ4751 | pGAL1-REH1 URA3 CEN Ampr | This work |
| pAJ4752 | pGAL1-REH1+10 URA3 CEN Ampr | This work |
| pAJ4763 | pGAL1-3xmyc-REH1 (WT) URA3 CEN Ampr | This work |
| pAJ4764 | pGAL1-3xmyc-REH1+6 URA3 CEN Ampr | This work |
| pAJ5501 | pGAL1-3xmyc-REHΔ6 URA3 CEN Ampr | This work |
| pAJ5503 | pGAL1-3xmyc-REH1Δ50 URA3 CEN Ampr | This work |

**Table II.**
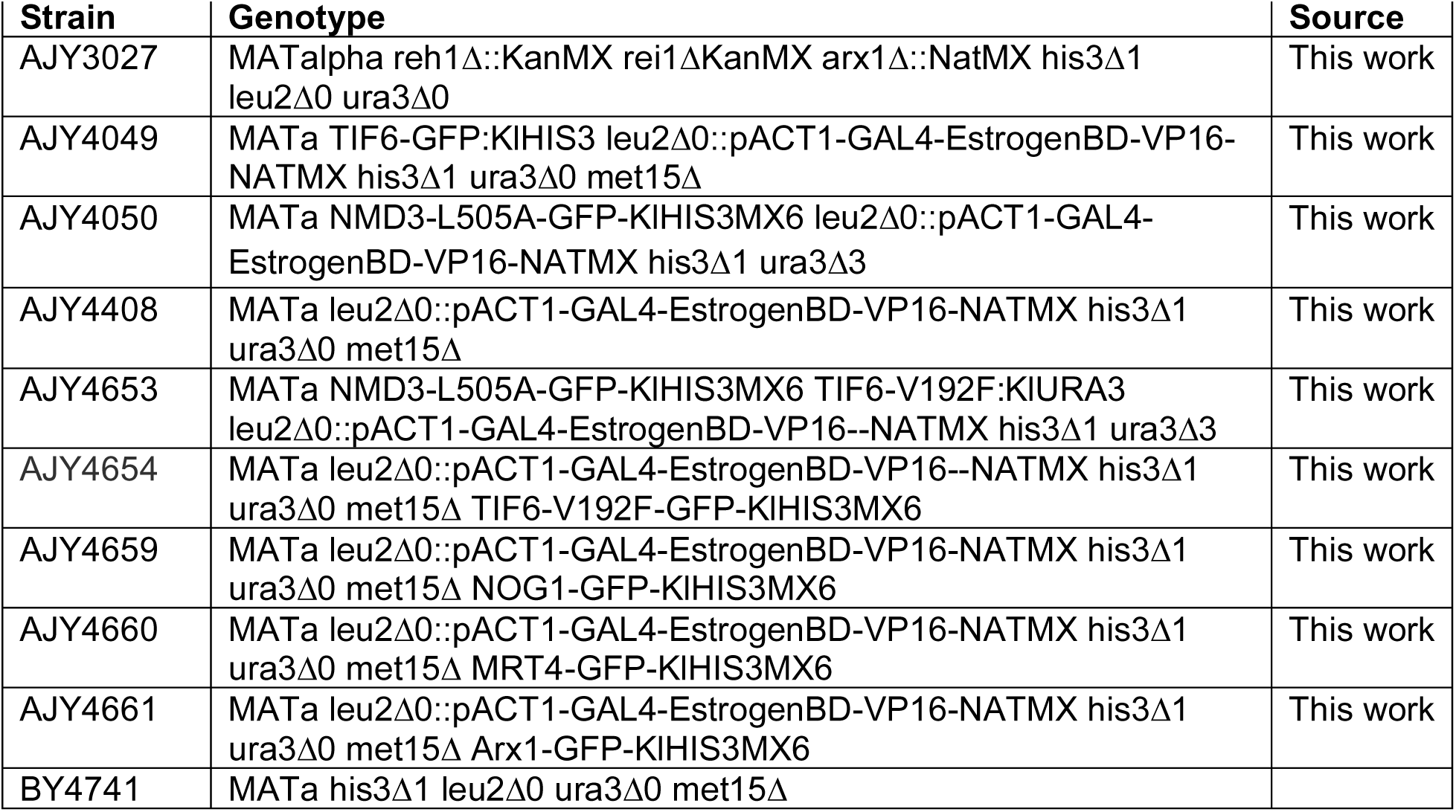
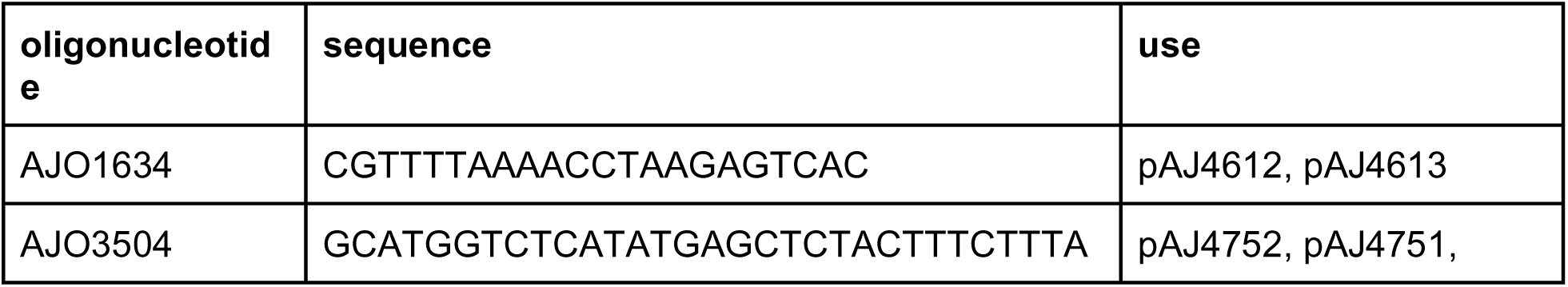
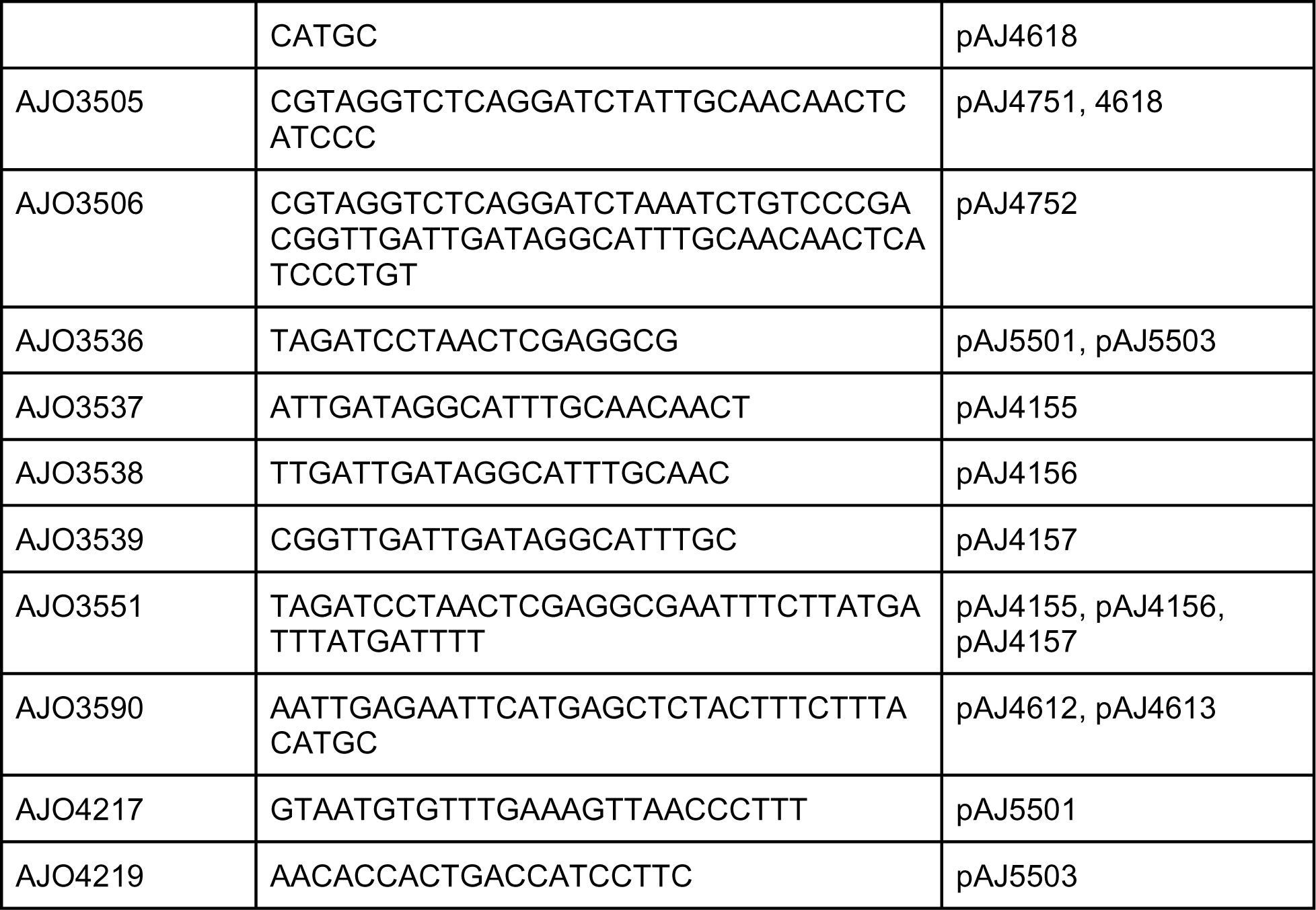
Strains.

## Materials and Methods

### Strains and Plasmids

#### Strains

AJY4049 was made by integrating PmeI-cut pAOL47-04 into a TIF6-GFP-expressing strain (Invitrogen). AJY3027 was constructed by sequential crossing appropriate haploid strains originally derived from the heterozygous deletion collection (Invitrogen). The arx1 deletion allele was converted from KanMX to NatMX by homologous replacement using p4339^39^. AJY4050 was made by integrating PmeI-cut pAOL47-04 into AJY4078 which in turn was made by integrating an NMD3-L505A-GFP HIS3MX6 cassette from pAJ3590. AJY4408 was made by integrating PmeI-cut pAOL47-04 into BY4741. Strains AJY4659, AJY4660 and AJY4661 were made by integrating PmeI-cut pAOL47-04 into the respective GFP strains (Invitrogen). AJY4654 was made by integrating PmeI-cut pAOL47-04 into AJY4047, which in turn was made by amplifying the TIF6-V192F-GFP::HIS3MX locus from AJY3941^38^ and integrating into BY4742. AJY4653 was made by integrating a TIF6-V192F:KlURA3 cassette into AJY4050.

#### Plasmids

pAJ4155, pAJ4156 and pAJ4157 were made from pAJ4752 by inverse PCR using AJO3537, AJO3538 and AJO3539, respectively, and the common oligo AJO3551. pAJ4751 and pAJ4157 were amplified with AJO3590 and AJO1634 and the amplicons digested with EcoRI and XhoI and cloned into the EcoRI and SalI sites of pMALC2H10T to make pAJ4612 and pAJ4613, respectively. pAJ4618 REH1 was amplified from genomic DNA using AJO3504 and AJO3505 and assembled via Golden Gate cloning with the appropriate part vectors from the Yeast Toolkit^40^. Yeast Toolkit plasmids were a gift from John Dueber (Addgene kit # 1000000061). pAJ4619 was made from pAJ4618 by inverse PCR using oligos AJO3551 and AJO3539. pAJ4621 was made by cloning a fragment containing P_GAL1_-ProtA-TEV-FLAG into the XbaI-SacI sites of pAJ4618. pAJ4751 REH1 was amplified from genomic DNA using AJO3504 and AJO3505 and assembled via Golden Gate cloning with appropriate Yeast Toolkit parts. pAJ4752: REH1 was amplified from genomic DNA using AJO3504 and AJO3506 and assembled with the appropriate Yeast Toolkit parts. pAJ4763 was made by moving the EcoRI-BglII pGAL1 fragment from pAJ4751 into same sites of pAJ4753. pAJ4753 was made by cloning a gene block (IDT) containing 500 nucleotides upstream of the REH1 start codon followed by a 3xmyc tag into the XbaI and SacI sites of pAJ4751. pAJ4764 was made by cloning a EcoRI-BglII pGAL1 fragment from pAJ4751 into same sites of pAJ4754 was made by cloning a gene block (IDT) containing 500 nucleotides upstream of the REH1 start codon followed by a 3xmyc tag into the XbaI and SacI sites of pAJ4157. pAJ5501 and pAJ5503 were made by inverse PCR using pAJ4763 and oligonucleotides AJO3536 with AJO 4217 and AJO4219, respectively. TIF6(V192F) was moved as a SacI, XhoI fragment from pRS316-TIF6(V192F) (A Warren) into pRS415 to make pAJ2249.

### Protein purification

#### Reh1

Reh1 was expressed and purified from E coli as an MBP-fusion. BL21 (DE3) codon+ cells (Agilent) carrying pAJ4612 (WT Reh1) or pAJ4613 (Reh1+6) were cultured in LB medium containing 0.2% glucose, 75 ug/ml ampicillin and 25 ug/ml chloramphenicol. When cells reached OD600=0.4, cells were shifted to 16°C and expression was induced with 1 mM IPTG overnight. All subsequent steps were performed at 0-4°C. Cells were washed and resuspended in lysis buffer (40 mM Tris-HCl, pH 8.0, 500 mM NaCl, 10% glycerol, 5 mM BME, 1mM PMSF and 1 uM each leupeptin and pepstatin). Cells were broken by sonication and the extract clarified by centrifugation at 25,000g for 20 min. Imidazole was added to 10 mM and protein was bound to Ni-NTA resin (Invitrogen) in bulk for 1 hr. Resin was transferred to a disposable column and washed with 15 column volumes of lysis buffer supplemented with 10 mM imidazole, followed by 15 column volumes of low salt buffer (lysis buffer with NaCl reduced to 50 mM) supplemented with 20 mM imidazole. Protein was eluted with low salt buffer supplemented with 250 mM imidazole. Peak fractions were pooled, supplemented with 1 mM EDTA and 1 mM DTT and bound to amylose resin. The resin was washed with 10 column volumes of amylose column buffer (40 mM Tris-HCl, pH 8.0, 500 mM NaCl, 10% glycerol, 1 mM EDTA, 1 mM DTT, 1 mM PMSF and 1 uM each leupeptin and pepstatin). Protein was eluted in amylose column buffer supplemented with 10 mM maltose. Peak fractions were pooled and applied to a Superose 12 column (Cytiva) equilibrated in amylose column buffer. Peak fractions were pooled and concentrated using a 30,000 mwco utrafiltration device (Amicon). Efl1, Sdo1 and 60S subunits were purified as described previously (Patchett). All protein concentrations were determined using a Qubit4 (Invitrogen).

### GTPase assays

Reactions (25ul) were set up on ice with amounts of Efl1, Sdo1, Nmd3, Reh1 (WT or mutant) and 60S specified in the figure legend in reaction buffer (20mM HEPES-KOH pH 7.5, 50mM KCl, 2mM Mg CH_3_COOH, 1mM DTT). Reactions were initiated by the addition of GTP to a final concentration of 40 uM containing a trace amount of [alpha-^32^P]-GTP (Perkin Elmer) and incubated for 20 min at 30°C. Reactions were quenched on ice with 6.25ul 0.5M EDTA and 1ul of each reaction was spotted onto a PEI-cellulose thin layer chromatography plate (Sigma-Aldrich) and developed in 0.8M CH_3_COOH and 0.8M LiCl. GTP and GDP were imaged with a storage phosphor screen on a Typhoon Phosphoimager. Data was analyzed using ImageJ software (NIH). All samples were corrected for non-enzymatic background hydrolysis.

### Sucrose density gradient sedimentation

Cultures were grown to OD600 of 0.3 in 250ml of selective media by continuous overnight growth. Where indicated, β-estradiol was added to a final concentration of 1uM and growth was continued for an additional 90-120 minutes. Cycloheximide (CHX) was added to a final concentration of 100µg/ml, cultures shaken for a further 10 minutes at 30°C before harvesting and storing at - 80°C. Cell pellets were washed and resuspended in 300ul of lysis buffer (20 mM Tris·HCl, pH 7.5, 100 mM NaCl, 30 mM MgCl2, 100 μg/ml cycloheximide, 200 μg/ml heparin, 5mM β-mercaptoethanol, 1mM PMSF and 1µM each of leupeptin and pepstatin). Extracts were prepared by agitation with glass bead and clarified by centrifugation at 4°C for 15 minutes at 18,000g. 10 A260 units of clarified extract were loaded onto 7-47% (w/v) sucrose gradients (containing 50 mM Tris·HCl, pH 7.5, 50 mM KCl, 12 mM MgCl2, 1 mM dithiothreitol [DTT]) and centrifuged for 2.5 hours at 40,000 rpm in a Beckman SW40 rotor. Gradients were fractionated using an ISCO Model 640 into 600µl fractions with continuous monitoring at 254nm. 1.2 ml 100% ethanol was added to each fraction, vortexed and stored at -20°C overnight. Fractions were centrifuged at 4°C for 15 minutes at 18,000g and pellets were dissolved in 1X SDS-PAGE sample buffer and heated at 99°C for 3 mins. Proteins were separated on 6-18% gradient SDS-PAGE gels, transferred to nitrocellulose membrane and subjected to western blot analysis using anti-myc (Biolegend), anti-Rpl8 antisera.

### Fluorescence Microscopy

The fluorescence signal was captured using a 500-ms exposure time on a Nikon E800 microscope fitted with a Plan Apo 100/1.4-numerical-aperture objective and a PCO sCMOS pco.edge camera controlled by NIS-Elements AR2.10 software, and photos were processed with Affinity Designer.

### Ribosome Profiling Experiments and Computational Analyses

Cultures of AJY4408 with plasmid pAJ4761 were grown to OD600 of 0.3 in 1.5 L of Leu-Glucose media by continuous overnight growth. β-estradiol was added to a final concentration of 1 uM and growth was continued for an additional 90 min. Cycloheximide (CHX) was added to a final concentration of 100 µg/ml and cultures were shaken for 10 minutes at 30°C, harvested and frozen at -80°C. Cells were washed once in IP buffer (Tris HCl 20 mM pH 7.5, 100 mM KCl, 10 mM MgCl2, 100 ug/ml CHX, Pierce Protease Inhibitor Cocktail, EDTA-free (1 tablet/10 ml), 1 mM PMSF, 1 uM each leupeptin and pepstatin and 5 mM beta-mercaptoethanol (BME)) and then resuspended in a volume of IP buffer equal to that of the cell pellet. Extracts were prepared by agitation with glass beads, Igepal was added to 0.1% extracts were clarified by centrifugation at 4°C for 15 minutes at 18,000 g. 20 mg of RNA (∼2 ml sample) was digested with 62.5 units of RNase I (Epicentre) at 4°C for 1 hr. 50ul was removed for total Ribo-seq sample. 25 ul bed volume washed anti-FLAG magnetic beads (Sigma Milipore) was added to the remaining sample and samples were rotated for 1 hr at 4°C. Beads were washed 4 times with IP buffer supplemented with 0.1% Igepal and transferred to a clean tube during the last wash. Reh1-bound ribosomes were eluted in 100 ul of IP buffer supplemented with 0.1% Igepal and 150 ug/ml 3xFLAG peptide (Sigma Milipore) for 15 min at 4°C. 12.5 units RNase I was added to the eluate and digestion was carried out at room temperature for 45 min. RNAse I digestion reaction was stopped by addition of QIAzol followed by RNA extraction by chloroform and ethanol precipitation.

For total RIbo-seq samples, 6.25 units RNase I (Epicentre) was added to 5ul of sample in a total volume of 50 ul in IP buffer supplemented with 0.1% Igepal. Digestion was carried out at room temperature for 45 min. Digested lysates were layered onto 50 ul 1M sucrose cushions prepared in IP buffer and ribosomes were pelleted by centrifugation at 70,000rpm in a TLA 100 rotor (Beckman Coulter) for 30 min at 4°C.

RNA was extracted from the ribosome pellets with the addition of 700 μl QIAzol followed by chloroform and ethanol precipitation. Isolated RNAs were size-selected using a 15% polyacrylamide TBE-UREA gel (Invitrogen) and 28-35 nt fragments were excised and extracted by crushing the gel fragment in 400 µL of RNA extraction buffer (300 mM NaOAc [pH 5.5], 5 mM MgCl_2_). The sample was passed through a Spin X filter (Corning 8160) and the flow through was ethanol precipitated (2.5 X volume) with 1.5 µL of Glycoblue (Invitrogen AM9516). Ribosome profiling libraries were prepared as previously described^41^ using the D-Plex Small RNA-Seq Kit (Diagenode). The cDNA was amplified for ten cycles. Each library was cleaned with PCR purification kit (Qiagen) and eluted with 30 μL of RNase-free water. To completely remove the primer-dimers, size-selection was done in a 3% agarose pre-cast gel (Sage Science) with the BluePippin system (Sage Science) using 180-235 nt range (tight settings, target:208 bp). The resulting size-selected libraries were analyzed with the Agilent High Sensitivity DNA Kit (Agilent). Each library was mixed in equimolar proportion for sequencing with the NovaSeq 6000 SP PE.

Ribosome profiling data analyses was preprocessed using RiboFlow^42^. Briefly, the first 12 nucleotides were extracted and used as unique molecular identifiers. The next four nucleotides matching the sequence ‘NGGG’ were discarded as these were added during the template-switching reverse transcription step. Sequencing fragments containing the 3’ adapter sequence AAAAAAAAAACAAAAAAAAAA were retained for further analyses. rRNA, tRNA and mRNA sequences were obtained from SGD on March 14, 2022. Sequences for the CDS (coding sequence), 5’ UTR (untranslated region) and 3’ UTR were merged to form the complete mRNA transcript giving us a total of 5024 transcripts. For transcripts annotated with more than one 3’ UTR sequence, we randomly retained one of the sequences. The resulting reference sequences are available at https://github.com/ribosomeprofiling/yeast_reference. PCR duplicates were eliminated from the ribosome profiling data using UMI-tools^43^. Statistical analyses and meta-gene plots were generated using RiboR using ribosome footprints of length between 26 and 29^44^.

### Northern blotting for tRNA enrichment

Cultures (500 ml) of yeast strain AJY4408 with plasmids pAJ4618 or pAJ4621 were grown and extracts prepared as described for Ribosome Profiling Experiments. Polysomes were collapsed to monosomes by digestion with 3 units of RNase I (Ambion) per A260 unit for 2 hr at 4°C. Immunoprecipitation was carried out by addition of ∼3 mg of magnetic Dynabeads coupled with rabbit IgG^45^ for 1 hr at 4°C. Beads were washed three times in IP buffer and particles eluted at 4°C with TEV protease (prepared in house). Eluted samples were diluted into LETS buffer (0.1 M LiCl, 0.01 M EDTA, 0.01 M Tris-Cl (pH 7.4) and 0.2% SDS), extracted sequentially with phenol/CHCl3 and CHCl3, and RNA was precipitated with ethanol with addition of Glycoblue (Ambion). Input RNA samples were isolated from 7.5 A260 units before or after RNaseI digestion, by phenol/CHCl3 extraction in LETS. Ribosome-bound RNA samples were prepared by sedimentation of 7.5 A260 units of RNaseI-treated extract through 50 ul of 17% sucrose in IP buffer for 20’ at 70,000 rpm in a Beckman TLA100 rotor at 4°C. Pellets were resuspended in LETS and RNA was extracted with phenol/CHCl3 followed by ethanol precipitation. For northern blotting, 5 ug of total RNA or 15% of the immunoprecipitated samples was separated on a 10% TBE-Urea gel (Invitrogen), transferred to Zeta-probe (BioRad) membrane and hybridized with the indicated radiolabeled oligonucleotide probes. Probes were: tRNA_i_ ^Met^ (5’-TGGTAGCGCCGCTCGGTTTCGAATCC), tRNA_e_^Met^ (5’-TGCTCCAGGGGAGGTTCGAACTCTCGACC) tRNA ^Arg^ (5’-CACTCACGATGGGGGTCGAA), and 5S (5’-TCTGGTAGATATGGCCGCAACC).

### Cryo-EM data collection and processing

Reh1-FLAG immunoprecipitation eluate was freshly prepared for cryo-electron microscopy. Quantifoil R1.2/1.3 grids coated with an ultrathin layer of amorphous carbon (Electron Microscopy Sciences) were glow-discharged for 1 minute at 25 mA. Using a Mark IV VitroBot (Thermo Fisher Scientific), 2.5 uL of sample was applied to freshly glow-discharged grids at 4 °C and 100% humidity, immediately blotted for 2 seconds at a force setting of 0, then plunged into liquid ethane and stored under liquid nitrogen until data collection. Microscopy data were collected at the University of Texas at Austin Sauer Structural Biology Center (RRID:SCR_022951), on a Titan Krios microscope (Thermo Fisher Scientific) operating at 300 kV, equipped with a K3 Summit direct electron detector (Gatan). Movies were acquired using SerialEM^46^ and recorded as 20 frames over 4 seconds for a total electron dosage of ∼70 e^-^ per Å^2^. Movies were recorded at a nominal magnification of 22,500x, a pixel size of 0.81 Å, and over a defocus range of -1 µm to - 2.5 µm. Using cryoSPARC Live for on-the-fly processing, movies were motion-corrected and dose-weighted, and CTF estimates performed. Particles of diameter between 300 Å and 400 Å were selected using a circular blob picker. On-the-fly 2D classification was used to separate particles in clean 80S and pre-60S classes, which were then exported to cryoSPARC (version 3.2), where multiple subsequent rounds of 2D classification resulted in a total of 217,663 particles corresponding to 80S and pre-60S projections. These particles were used for *ab initio* reconstruction and 3D heterogeneous refinement into separate 80S and pre-60S classes of 115,111 and 92,218 particles, respectively.

The pre-60S class showed complete or partial occupancy for cytoplasmic ribosome assembly factors, including Tif6, Nmd3, Lsg1 and Reh1, but further analysis was precluded by severe orientation bias. The well-resolved 80S class was further subjected to a two-part 3D classification scheme in cryoSPARC, with a soft mask surrounding the A site followed by a soft mask surrounding the E site. This scheme led to four 80S classes containing A and P site tRNAs, and/or E site eIF5A (Supplementary Figure S5B).

### Modeling, refinement and graphics

PDB 7NRC^47^ was first rigid-body docked into the two maps with resolved tRNAs, with and without eIF5A. The model for Reh1 was derived from PDB 6QTZ^5^. A predicted model for yeast Rps25-A (AF-Q3E792-F1) was downloaded from the AlphaFold Protein Structure Database^48,49^, and its N-terminus was fit manually into the EM map using COOT^50^. The mRNA chain was built manually in COOT. Models were relaxed using flexible fitting implemented in ISOLDE^51^, then subjected to real-space refinement implemented in PHENIX^52^. Molecular graphics were prepared using UCSF ChimeraX^53^.

**Figure 3 S1.**
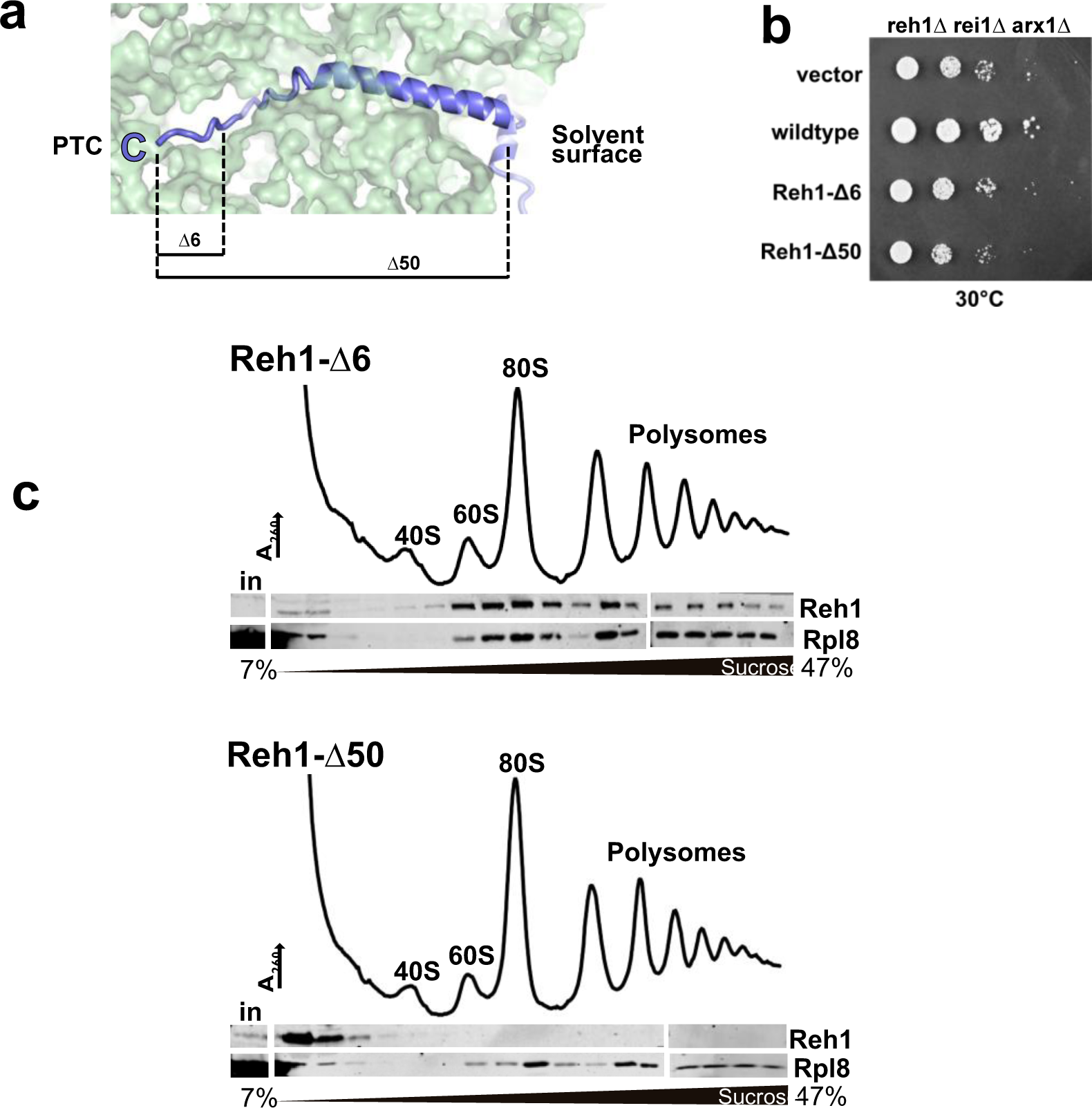
The C-terminus of Reh1 is necessary for function and contributes to ribosome binding. *Panel A:* Cartoon of the C-terminus of Reh1 extended along the length of the polypeptide exit tunnel. Endpoints of Reh1Δ6 and Reh1Δ50 C-terminal truncations are indicated. *Panel B:* Complementation assay for REH1 variants. Ten-fold serial dilutions of rei1Δ arx1Δ reh1Δ mutant cells (AJY3027) containing vector or vectors expressing REH1 WT, reh1Δ6 or reh1Δ50 grown on Ura-medium. *Panel C:* Sedimentation of Reh1Δ6. *Panel D:* Sedimentation of Reh1Δ50

**Figure 3 S1.**
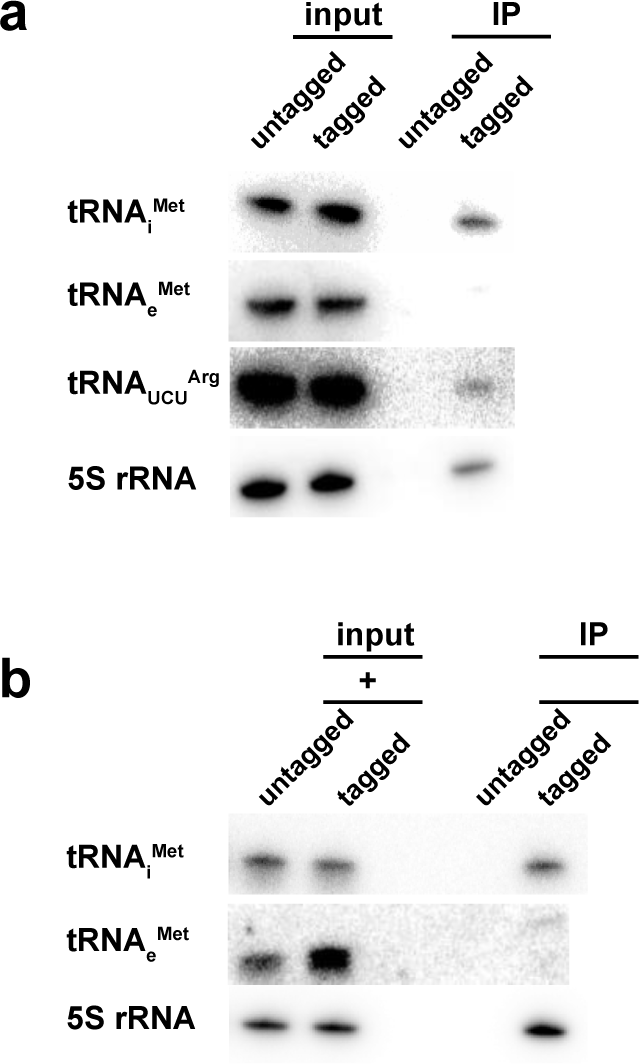
Reh1-containing 80S are enriched for tRNA_i_^Met^. *Panel A:* As described for Figure 3, except input was total RNA before RNaseI digestion. *Panel B:* As described for Figure 3, except input was total RNA after RNaseI digestion.

**Figure 4 S1.**
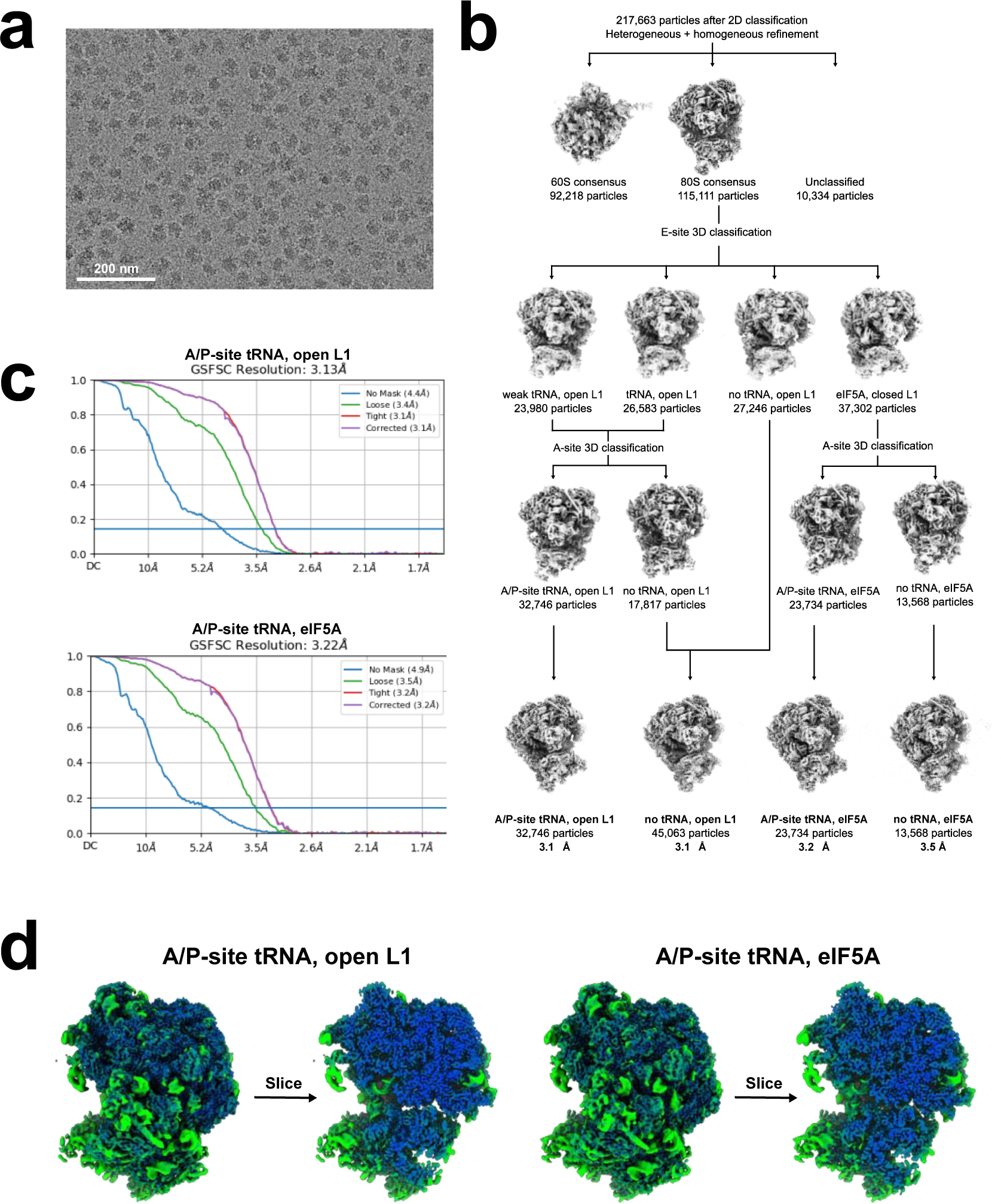
Cryo-EM data processing and validation. *Panel A:* Representative micrograph of vitrified Reh1-80S sample. *Panel B:* Cryo-EM data processing workflow. *Panel C:* Gold-standard Fourier shell correlations (GSFSCs) for the two tRNA-containing Reh1-80S structures. Standard cutoff = 0.143. *Panel D:* Local resolution analysis of the two tRNA-containing Reh1-80S structures.

**Figure 5 S2.**
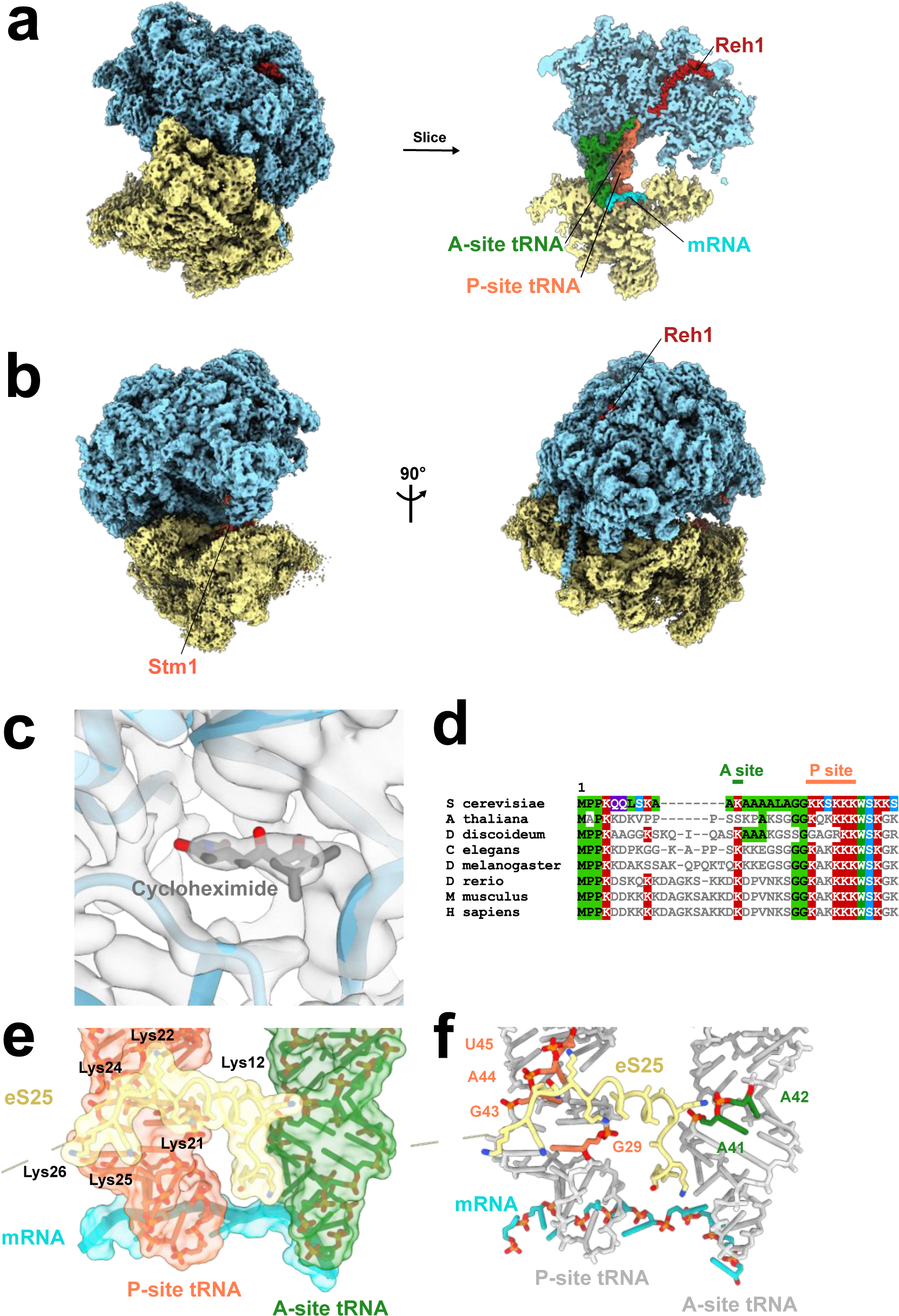
Additional cryo-EM structural data. *Panel A:* Cryo-EM map (left) and cross-section (right) of the Reh1-80S structure with Reh1 (red), A site tRNA (green), P site tRNA (orange) and mRNA (turquoise). *Panel B:* Cryo-EM map of the Reh1-80S structure without tRNAs and without eIF5A, with partial occupancy of the ribosome dormancy factor Stm1 (orange). *Panel C:* Cryo-EM map of the E site, with occupancy of cycloheximide shown. *Panel D:* Multiple sequence alignment of eS25 from various eukaryotic species. Identical matches to the top sequence (yeast) are shown with a colored background. Residues interacting with yeast A and P site tRNAs are indicated at the top. *Panel E:* Surface (left) and skeletal (right) interactions between the N-terminus of eS25 and peptidyl transferase center (PTC) ligands in the structure of Reh1-80S-tRNA. Left, lysines of yeast eS25 interacting with tRNA are indicated. Right, tRNA nucleotides interacting with eS25 are indicated.

